# Genomic diversity and antimicrobial resistance of *Staphylococcus aureus* in Saudi Arabia: a nationwide study using whole-genome sequencing

**DOI:** 10.1101/2025.03.03.641358

**Authors:** Mohammed S. Alarawi, Musaad Altammami, Mohammed Abutarboush, Maxat Kulmanov, Dalal M. Alkuraithy, Senay Kafkas, Robert Radley, Marwa Abdelhakim, Hind Masfer Abdullah Aldakhil, Reema A. Bawazeer, Mohammed A. Alolayan, Basel M. Alnafjan, Abdulaziz A. Huraysi, Amani Almaabadi, Bandar A Suliman, Areej G Aljohani, Hassan A. Hemeg, Turki S Abujamel, Anwar M. Hashem, Ibrahim A Al-Zahrani, Mohammed S Abdoh, Haya I Hobani, Rakan F Felemban, Wafaa A Alhazmi, Pei-Ying Hong, Majed F. Alghoribi, Sameera Aljohani, Hanan Balkhy, Abdulrahman Alswaji, Maha Alzayer, Bassam Alalwan, Mai M. Kaaki, Sharif M. Hala, Omniya Ahmad Fallatah, Wesam Ahmad Bahitham, Samer Yahya Zakri, Mohammad A Alshehri, Nader Kameli, Abdullah Algaissi, Edrous Alamer, Abdulaziz Alhazmi, Amjad A.Shajri, Majid Ahmed Darraj, Bandar Kameli, O. O. Sufyani, Badreldin Rahama, Abrar A. Bakr, Fahad M. Alhoshani, Azzam A. Alquait, Ali Somily, Ahmed M. Albarrag, Lamia Alosaimi, Sumayh A. Aldakeel, Fayez S. Bahwerth, Tamir T Abdelrahman, Séamus Fanning, Essam J. Alyamani, Takashi Gojobori, Satoru Miyazaki, Mohammed B. Al-Fageeh, Robert Hoehndorf

## Abstract

Methicillin-resistant *Staphylococcus aureus* (MRSA) surveillance in regions with mass gatherings presents unique challenges for public health systems. Saudi Arabia, hosting millions of pilgrims annually, provides a distinctive setting for studying how human mobility shapes bacterial populations, yet comprehensive genomic surveillance data from this region remains limited. Here, we present an integrated analysis of *S. aureus* isolates collected across seven Saudi Arabian regions, combining whole-genome sequencing with extensive antimicrobial susceptibility testing and standardized metadata following FAIR data principles. Our analysis revealed striking differences between pilgrimage and non-pilgrimage cities. Pilgrimage cities showed significantly higher genetic diversity and antimicrobial resistance rates, harboring numerous international strains including recognized clones from diverse geographic origins. Reported lineage dynamics is changing, expanding toward community clones. While genomic prediction of antimicrobial resistance showed high accuracy for some antibiotics, particularly beta-lactams, with varying performance for others, highlighting the necessity for phenotypic testing in clinical settings. Our findings demonstrate how mass gatherings drive bacterial population structures and emphasize the importance of integrated surveillance approaches in regions with significant global connectivity and travel.

**Importance:** Genomic data enables the tracking of pathogens by revealing clonal expansions within populations and identifying successful lineages. However, comprehensive national-level data from Saudi Arabia remains limited on a large scale. The adoption of FAIR principles and reproducible workflows ensures robust, consistent analysis, fostering effective data sharing. The OneHealth approach’s success depends on the integration and collaboration across diverse domains in today’s digital landscape.

## Introduction

*Staphylococcus aureus*, a Gram-positive bacterium characterized by its grape-like clustering morphology, is both a common human and animal commensal organism and a significant pathogen that has emerged as a leading cause of hospital-acquired infections. Notably, methicillin-resistant *S. aureus* (MRSA) accounts for approximately 25-50% of these infections. MRSA emerged shortly after methicillin’s introduction in 1958, following the acquisition of a mobile genetic element known as the staphylococcal chromosomal cassette `mec` (`SCCmec`). This cassette harbors the mec*A* gene, which encodes PBP2A, an alternative penicillin-binding protein involved in cell-wall synthesis. The modified protein exhibits reduced affinity to *β*-lactam antibiotics, conferring broad-spectrum resistance to this antibiotic class [30, 32].

The global impact of MRSA has expanded significantly beyond healthcare settings, establishing distinct reservoirs within communities and livestock populations [34, 21]. Understanding the epidemiological distribution of MRSA lineages and their corresponding resistance phenotypes across communities, regions, and countries has become increasingly crucial from public health prospective. This knowledge directly informs the development of targeted mitigation strategies, encompassing evidence-based policies for antibiotic stewardship, systematic screening protocols, and comprehensive livestock management practices [5, 42]. The clinical significance of MRSA is particularly evident in its role as a major pathogen in post-surgical complications, community-onset infections, and foodborne illness outbreaks [34, 6].

Saudi Arabia presents a unique context for studying MRSA transmission and resistance patterns. As the largest country on the Arabian Peninsula by both area and population, it holds profound religious significance as the custodian of Islam’s two holiest sites. This unique position draws millions of Muslim pilgrims annually to the Kingdom. Mass gatherings increase chances for microbial transmission and the introduction of novel clones and antimicrobial resistance patterns. Large cities are hypothesized to function as entry points and subsequent dissemination of imported clones due to population density and connectivity. Consequently, international clones are predicted to emerge initially in these entry points before spreading in other regions within Saudi Arabia. Smaller, less connected cities are expected to display a less presence of these clones. The epidemiology of MRSA in Saudi Arabia has been documented through various investigations [6, 47, 2, 14], although most studies have been limited to individual healthcare facilities or specific geographical regions. These investigations range from comprehensive phenotypic characterization of MRSA isolates’ antimicrobial susceptibility profiles [2, 7, 1] to more detailed molecular analyses of strain characteristics [47].

Despite extensive research on MRSA in Saudi Arabia, there remains a significant gap in comprehensive datasets that integrate both genotypic and phenotypic information across multiple geographical regions. One critical component to address this gap is the use of FAIR (Findable, Accessible, Interoperable, Reusable) data principles [54]. Implementation of FAIR increases the value of accumulated data by enabling easier reuse and interoperability. Interoperable data is valuable because it can be combined with other datasets and aid in elucidating molecular mechanisms of antimicrobial resistance and identifying novel therapeutic targets in addition to transmission burden. Moreover, combined genotype–phenotype datasets enable the development of machine learning approaches for predicting drug resistance patterns and pathogenicity profiles, while also informing the design of targeted studies aligned with the One Health approach.

To address this knowledge gap, we established a collection of 686 *S. aureus* isolates from diverse regions across Saudi Arabia, encompassing clinical specimens, community screening samples, and wastewater isolates. We conducted whole-genome sequencing on the isolates and performed extensive drug resistance phenotyping screening, creating a novel geospatial genotype–phenotype dataset. Our analysis reveals high genetic diversity, including several previously unidentified sequence types distributed across different regions. These novel sequence types exhibit elevated patterns of drug resistance. Significantly, we observed higher genetic diversity and increased levels of drug resistance in cities associated with mass gatherings (Jeddah, Makkah, Madinah), suggesting that mass gathering may serve as drivers of antimicrobial resistance evolution and transmission. These findings provide crucial insights for developing targeted intervention strategies and underscore the importance of genomic surveillance in regions experiencing regular mass gatherings.

## Materials and Methods

### Sample Selection and Sourcing

Our study includes a total of 686 *S. aureus* isolates collected from seven distinct regions across Saudi Arabia. The majority of samples were obtained from tertiary care hospitals, with the Western region contributing the largest proportion: Jeddah (275 isolates), Madinah (102 isolates), and Makkah (36 isolates). Additional regional collections include Riyadh in the Central region (158 isolates), Hail in the North (64 isolates), AlHasaa in the East (33 isolates), and Jazan in the South (18 isolates). To broaden the scope of our surveillance, we supplemented the hospital-sourced isolates with 60 samples collected through a community screening of healthy individuals in Jeddah city and 5 environmental wastewater samples from the same region. A detailed breakdown of sample distribution is provided in supplementary Table S1. Sample selection within each hospital followed a random sampling approach, and we included both methicillin-resistant (MRSA) and methicillin-susceptible *S. aureus* (MSSA) isolates to ensure comprehensive representation of circulating strains.

### Bacterial Identification and Susceptibility Testing

All isolates were initially enriched on tryptic soy agar (TSA) supplemented with 5% [w/v] NaCl (Sigma-Aldrich, Germany) to selectively culture *S. aureus*. Single colonies were used for subsequent bacterial identification, susceptibility testing, and glycerol stock archival. Throughout the identification process, the *S. aureus* ATCC 29213 strain served as an internal reference control. Phenotypic drug resistance profiling was performed using the VITEK^®^ 2 System with GP ID cards for identification and AST-P580 cards for antimicrobial susceptibility testing (bioMérieux, France). Supplementary table S2 shows the antimicrobial agents and their concentrations of the AST-P580 card. All test results were digitally recorded and stored for analysis.

### DNA Isolation

Genomic DNA isolation was performed using single colonies obtained either directly from Petri dishes or from enriched broth cultures following identification. For each sample, 2 mL of overnight culture was pelleted for DNA extraction. Concurrently, we prepared archived samples by combining 500 µL molecular grade glycerol (Thermofisher, USA) with 500 µL of the culture for future investigations. Genomic DNA was isolated using the automated KingFisher MagMax DNA isolation kit protocol (Thermofisher, USA). DNA quality and quantity were assessed using both a nano-drop instrument (Thermofisher, USA) and Qubit dsDNA BR kit (Thermofisher, USA).

### Library Preparation and DNA Sequencing

Genomic DNA libraries were constructed using 100 ng of input DNA with the QIASeq FX DNA library kit (Qiagen, Germany) following the manufacturer’s protocol. Library quality was assessed using an Agilent Bioanalyzer system (Agilent, USA). Whole-genome sequencing was performed on the Illumina NovaSeq 6000 platform (Illumina, USA) using SP flow-cells, generating 150-base paired-end reads with average insert sizes ranging from 350 to 450 bp, and 50-100X sequence depth.

### Genomic Analysis

Initial quality control and preprocessing of raw sequencing reads were perfomred using TrimGalore v0.4.4 for adapter trimming [9]. Taxonomic identification and contamination utilized Kraken v2.0.8 beta [55] and Mash v2.3 [39]. Genomes were assembled using SKESA v2.4.0 [49] and called variants using Snippy v4.6.0 against the reference genome `NC_007795.1`.

Annotation of the assembled genomes utilized prokka v1.14.6 [46] and Roary for pangenome construction v3.13.0 [40]. A neighbor-joining phylogenetic tree was built using FastTree2 2.1.10 [43] based on the core genome alignment. Assignment of multilocus sequence types (MLST) performed using ABRicate v1.0.1 with the MLST tool.

Multiple approaches were employed for antimicrobial resistance analysis. Drug resistance genes were identified using both the CARD v5.1.1 [3] and ResFinder v4.0 [18] databases via ABRicate v1.0.1 [52]. Additionally, DeepARG v1.0.2 [10], a deep learning approach, to enhance resistance gene detection. Virulence factors were identified using the Virulence Factor Database (VFDB) [22] through ABRicate v1.0.1. Methicillin resistance mechanisms were characterized by screening for the mec*A* gene using the staphopia-Sccmec v1.0.0 typing tool [41]. The analysis was complemented using a local database containing whole cassette sequences (mec*A*, mec*B*, mec*C*), cassette recombinase genes (CCR) genes, and insertion sequences for each SCC*mec* type. Additional mec*A* cassette detection was performed using minimap2 v2.24-r1122 [35] with a local database. The SCC*mec* type assignments were determined based on cassette element arrangements following the workflow at https://github.com/cdnstp/SCCmec_CLA. Finally, Protein A gene polymorphism (spa) typing was performed using spaTyper v4.6.1 [44] with its associated database.

### Reproducibility and Implementation of FAIR Data Principles

We implemented our entire analysis workflow using the Common Workflow Language (CWL) [23] to ensure computational reproducibility. The two main analysis workflows are shown in Supplementary Figures S1 and S2.

We executed all analyses on a local Arvados system [The Arvados Authors et al.], with complete workflow definitions and execution environments preserved. To promote Findability, Accessibility, Interoperability, and Reusability (FAIR), we have made all source code, workflow definitions, and configuration files freely available at https://github.com/bio-ontology-research-group/mrsa-sequences. The repository includes detailed documentation of computational requirements, software versions, and usage instructions. Additionally, raw sequencing data were deposited in the European Nucleotide Archive (ENA) under accession number `PRJEB59751`. Furthermore, the metadata of our samples as well as the measured antimicrobial resistance phenotypes using the Resource Description Framework (RDF) [33] are made available in a public repository [36] and on a Github repository. Supplementary Figure S3 describes the RDF data model.

## Results

### Genetic diversity and geographic distribution of *S. aureus* in Saudi Arabia

686 samples of *S. aureus* from multiple regions in Saudi Arabia were sequenced, following quality control validation, and confirmation as *S. aureus* through whole-genome sequencing and phenotypic testing. The isolates consist of methicillin-resistant (MRSA) and methicillin-susceptible (MSSA) strains, with MSSA proportions varying significantly by region: from 3.9% in Madinah to 34.4% in Hail (Table 1).

**Table 1:** Summary of MRSA statistics across different geolocations, including total counts, unique clonal complexes (CCs), unique sequence types (STs), SCC*mec* types rate, Panton-Valentine leukocidin (PVL) rate, and toxic shock syndrome toxin (TSST) rate.

| Geolocation | Total samples | Unique CCs | Unique ST | Sccmec types rate % | PVL rate % | TSST rate % |
| --- | --- | --- | --- | --- | --- | --- |
| Alhasa (East) | 33 | 8 | 15 | IVa 45.5; V 30.3; IVc 12.1; MSSA 6.1; VI 6.1 | 24.2 | 6 |
| Hail (North) | 64 | 9 | 19 | MSSA 34.4; IVa 28.1; V 23.4; VI 6.3; IVc 4.7; IIIa 3.1 | 6.3 | 26 |
| Jazan (South) | 18 | 4 | 6 | V 61.1; IVa 27.8; IVd 11.1 | 22.2 | 0 |
| Jeddah (West) | 275 | 9 | 43 | V 45.1; IVa 32.4; MSSA 10.2; IVc/IVd 3.6; VI 2.9; Undetermined 0.7; VI/Vc/IVh/IIIa 0.4 | 13.5 | 8 |
| Madina (West) | 102 | 9 | 26 | V 34.3; IVa 33.3; IVc 13.7; VI 6.9; MSSA 3.9; IVg 2.0; I 2.0; Others 1.0 | 32.4 | 13 |
| Makkah (West) | 36 | 8 | 14 | V 41.7; IVa 38.9; MSSA 13.9; Undetermined 2.8; IVc 2.8 | 13.9 | 8 |
| Riyadh (Central) | 158 | 9 | 29 | V 41.8; IVa 19.6; MSSA 18.4; IVc 8.9; IVd 4.4; VI 3.2; IVb 1.9; IVg/IVh 0.6 | 19.0 | 7 |

Molecular typing through PubMLST [31] classification was performed to understand the evolutionary relationships and population structure of these isolates, as this knowledge is crucial for tracking transmission patterns and implementing targeted infection control measures. The classification revealed nine major clonal complexes, with CC5 (26.38%), CC97 (12.39%), and CC22 (9.48%) being the most prevalent (Supplementary Figures S4). 31.92% of isolates belonged to previously unassigned clonal complexes. Our analysis also identified 14 novel sequence types (Supplementary Table S3). Figure 1 provides an overview of the molecular diversity and relatedness of the samples included in our study, in addition to genetics distance based on core Single Nucleotide Variants (SNVs) minimum spanning tree (Supplementary Figure S5).

**Figure 1:**
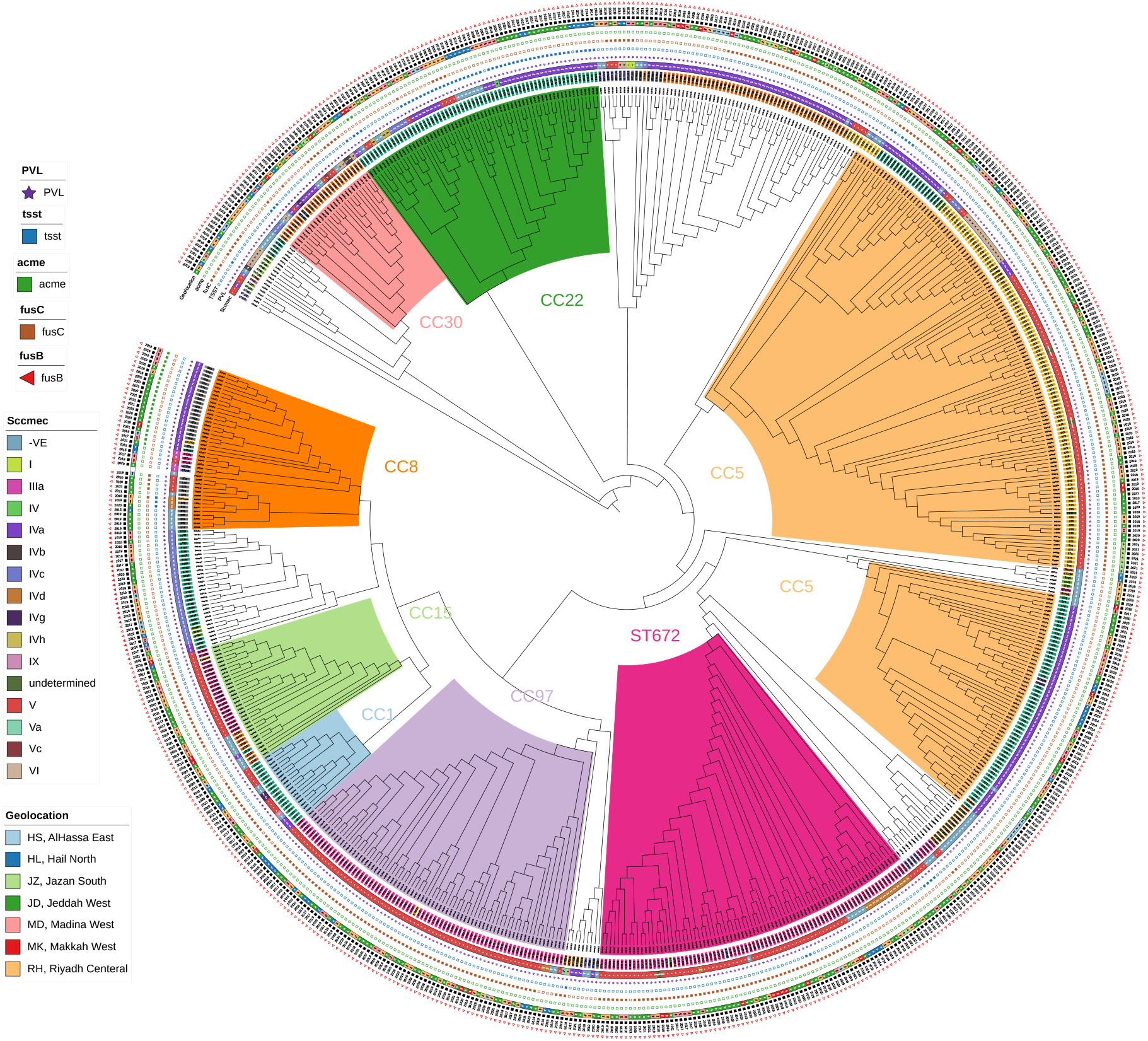
Phylogenomic structure and diversity of *S. aureus* across Saudi Arabia. Maximum likelihood tree constructed from core genome SNPs of 686 isolates. Major Clonal Complexes (CCs), grouping related Sequence Types, are highlighted (the dominant CC5, CC8, CC22, ST672). Coloured rings display the associated SCCmec type (inner ring) and geographical region of origin (outer ring), revealing the distribution patterns of major lineages. CC5 two branches (ST5, and ST6) and the distinct clustering reflecting national population structure influenced by regional and international factors. Key virulence/resistance markers (PVL, TSST, ACME, fusB/C) are indicated in the outermost tracks.

We observed distinct patterns of strain distribution aligned with pilgrimage routes in major cities. Cities along the Hajj and Umrah routes exhibited high genetic diversity: Jeddah (9 CCs, 43 STs), Madinah (9 CCs, 26 STs), and Makkah (8 CCs, 14 STs), likely reflecting the international convergence of travelers. In contrast, regions with less international traffic showed more limited diversity, particularly Jazan (4 CCs, 6 STs). Analysis of the SCC*mec* revealed region-specific patterns, with SCC*mec*V ranging from 23.4% in Hail to 61.1% in Jazan, and SCC*mec*IVa ranging from 19.6% in Riyadh to 45.5% in AlHasa (Table 1, Figure 2, and Supplementary Figure S6).

**Figure 2:**
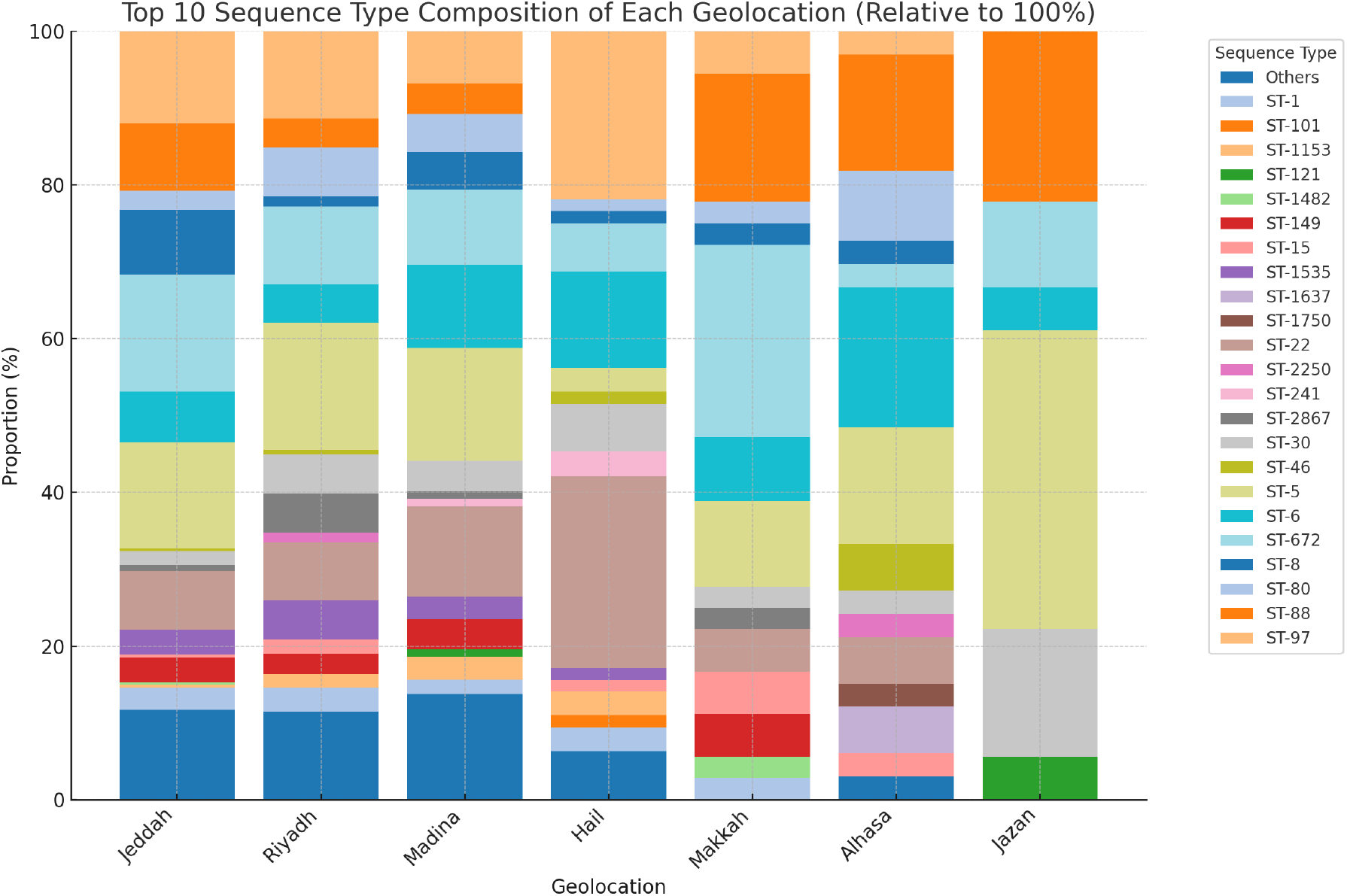
Stacked bar chart illustrating the relative proportion (%) of the top 10 most frequent *S. aureus* isolates Sequence types per region from Saudi Arabia. The chart highlights the greater ST diversity observed in major pilgrimage-associated cities (Jeddah, Madinah, Makkah) and other high-traffic locations like Riyadh, compared to regions like Jazan, which shows dominance by fewer STs (e.g., ST5, ST30). Notable STs like ST5, ST97, ST672, ST8, and ST22 show varying prevalence across regions, indicating complex population structures influenced by both local factors and international travel patterns regions,.

Several internationally recognized strains were observed in pilgrimage cities. The Bengal Bay clone (CC1-ST772-PVL positive) [38] and CC30-ST1482 [13] were detected primarily in Makkah and Jeddah, regions that host large numbers of international pilgrims. The South Pacific Clone (CC30-ST30) [53] was found across most cities.

### Phenotypic antimicrobial resistance

Antimicrobial susceptibility testing was performed on all isolates (Table 2). We observed the highest resistance rates for benzylpenicillin across all regions, ranging from 88.9% in AlHasa to 100% in Makkah. Oxacillin resistance, mediated by mec*A*, showed considerable variation, from 60.3% in Hail to 96.6% in Madinah. Fusidic acid emerged as the next most prevalent resistance, with particularly high rates in Riyadh (70.5%) and Jeddah (70.9%) but notably lower in Hail (35.3%). Fluoroquinolone resistance varied significantly by region, with levofloxacin resistance ranging from 14.7% in Hail to 45.7% in Madinah, and similar patterns for moxifloxacin (13.4% to 41.5%). Erythromycin resistance rates ranged from 19.4% in AlHasa to 37.9% in Madinah. We observed variable resistance to aminoglycosides, with tobramycin resistance ranging from 8.1% in Makkah to 32.5% in Jeddah, while gentamicin resistance showed similar patterns. Tetracycline resistance rates varied considerably, from 6.2% in AlHasa to 26.5% in Jeddah.

**Table 2:**
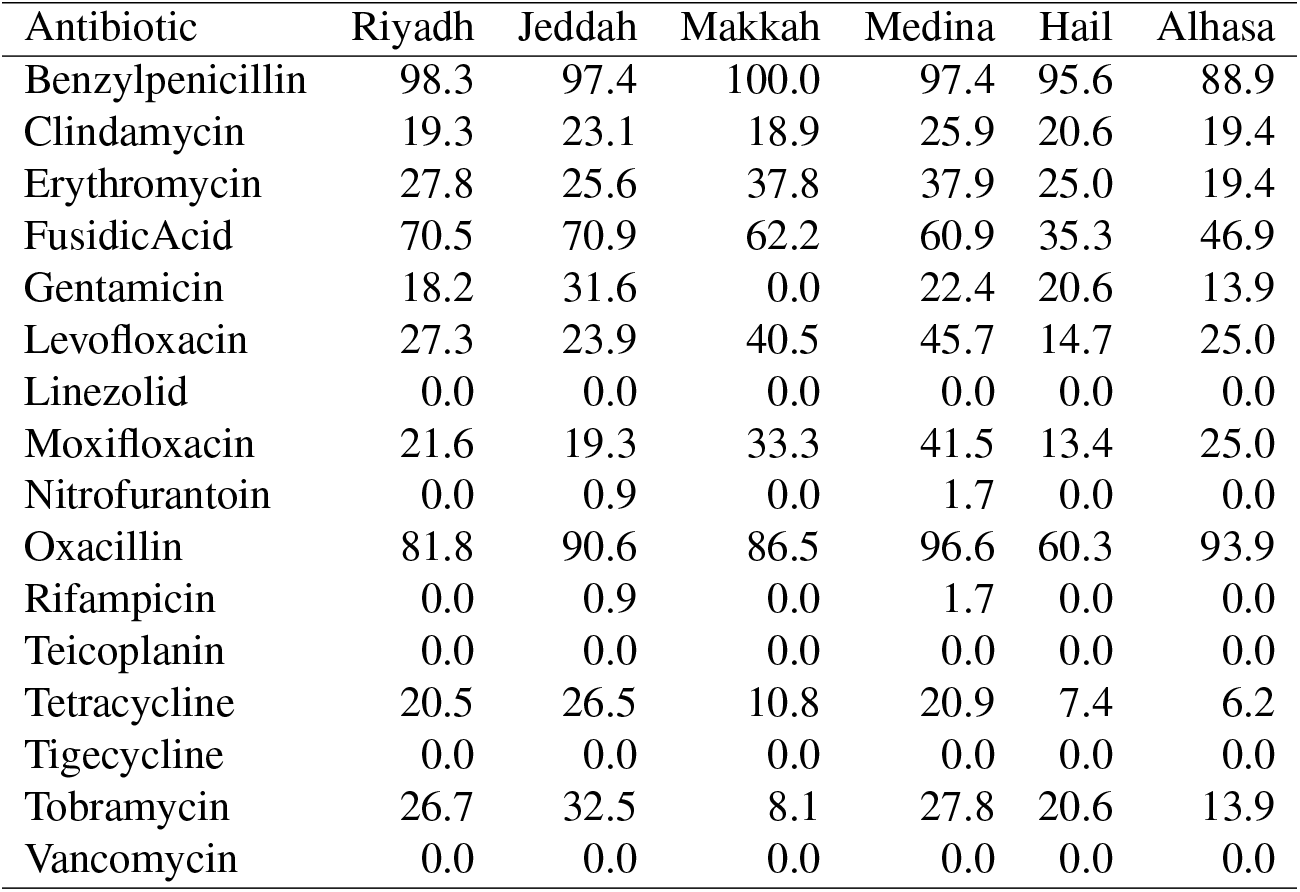
Antibiotic resistance rates by sampling location in percent.

| Antibiotic | Riyadh | Jeddah | Makkah | Medina | Hail | Alhasa |
| --- | --- | --- | --- | --- | --- | --- |
| Benzylpenicillin | 98.3 | 97.4 | 100.0 | 97.4 | 95.6 | 88.9 |
| Clindamycin | 19.3 | 23.1 | 18.9 | 25.9 | 20.6 | 19.4 |
| Erythromycin | 27.8 | 25.6 | 37.8 | 37.9 | 25.0 | 19.4 |
| FusidicAcid | 70.5 | 70.9 | 62.2 | 60.9 | 35.3 | 46.9 |
| Gentamicin | 18.2 | 31.6 | 0.0 | 22.4 | 20.6 | 13.9 |
| Levofloxacin | 27.3 | 23.9 | 40.5 | 45.7 | 14.7 | 25.0 |
| Linezolid | 0.0 | 0.0 | 0.0 | 0.0 | 0.0 | 0.0 |
| Moxifloxacin | 21.6 | 19.3 | 33.3 | 41.5 | 13.4 | 25.0 |
| Nitrofurantoin | 0.0 | 0.9 | 0.0 | 1.7 | 0.0 | 0.0 |
| Oxacillin | 81.8 | 90.6 | 86.5 | 96.6 | 60.3 | 93.9 |
| Rifampicin | 0.0 | 0.9 | 0.0 | 1.7 | 0.0 | 0.0 |
| Teicoplanin | 0.0 | 0.0 | 0.0 | 0.0 | 0.0 | 0.0 |
| Tetracycline | 20.5 | 26.5 | 10.8 | 20.9 | 7.4 | 6.2 |
| Tigecycline | 0.0 | 0.0 | 0.0 | 0.0 | 0.0 | 0.0 |
| Tobramycin | 26.7 | 32.5 | 8.1 | 27.8 | 20.6 | 13.9 |
| Vancomycin | 0.0 | 0.0 | 0.0 | 0.0 | 0.0 | 0.0 |

Several antibiotics maintained complete efficacy across all regions. We observed no resistance to linezolid, tigecycline, or teicoplanin. Vancomycin, often considered a last-resort antibiotic [17], showed nearly complete effectiveness, with only a single borderline case identified in a sample from Madinah with updated MIC break point. Similarly, nitrofurantoin and rifampicin maintained high effectiveness, with resistance rates not exceeding 1.7% in any region.

Comparison of overall differences of antimicrobial resistance was performed for locations associated with mass gatherings and other cities. Pilgrimage-associated cities (Madinah, Makkah, and Jeddah) showed significantly higher overall resistance rates compared to other locations (33.9% vs 27.6%, *p* = 0.047, one-tailed *t*-test).

### Genotypic identification of antimicrobial resistance

While whole-genome sequencing data identify resistance-associated genotypes, linking these directly to observed phenotypes remains a complex process. We compared phenotypic resistance results to genetic predictions using three approaches: CARD [4], ResFinder [18], and the deep learning tool DeepARG [10]. Genetic predictions showed varying levels of concordance with phenotypic resistance across different antibiotic classes (Table 3).

**Table 3:** Concordance of antimicrobial resistance predictions with the observed resistance patterns using three tools (CARD Database, Resfinder 4.0, and DeepARG). Metrics shown as Precision/Recall/F-score. Dash (-) indicates that no predictions could be made by the tool for this antimicrobial agent.

| Antibiotic | CARD | Resfinder | DeepARG | Concordance |
| --- | --- | --- | --- | --- |
| Benzylpenicillin | .983/.980/.982 | .994/.572/.726 | .983/.980/.982 | High CARD-DeepARG |
| Cefoxitin | 1.0/.004/.007 | .984/.950/.967 | - | Low |
| Oxacillin | .858/.976/.913 | - | .858/.976/.913 | High CARD-DeepARG |
| Gentamicin | .172/.937/.291 | .752/.793/.772 | .172/.937/.291 | Low |
| Tobramycin | .220/.950/.357 | .785/.807/.796 | .221/.950/.358 | Low CARD-DeepARG |
| Erythromycin | .283/1.0/.441 | .776/.889/.828 | .584/.895/.707 | Moderate |
| Clindamycin | .446/.861/.588 | .704/.826/.760 | .378/.861/.525 | Moderate |
| Tetracycline | .167/.962/.285 | .853/.771/.810 | .491/.800/.609 | Low |
| Fusidic Acid | .937/.925/.931 | .940/.845/.890 | 1.0/.005/.010 | High CARD-Resfinder |

Beta-lactam predictions demonstrated high accuracy, particularly for benzylpenicillin, where both CARD and DeepARG achieved identical high performance (precision 0.983, recall 0.980), while ResFinder showed higher precision (0.994) but lower recall (0.572). Oxacillin predictions showed strong agreement between CARD and DeepARG (precision 0.858, recall 0.976), while ResFinder did not provide predictions for this antibiotic. For fusidic acid, both CARD and ResFinder achieved accurate results (precision 0.937 and 0.940 respectively), while DeepARG could only make few predictions.

However, we observed substantial discrepancies for other antibiotic classes. Aminoglycoside resistance predictions varied markedly across tools. For gentamicin and tobramycin, CARD and DeepARG showed low precision (0.172–0.221) but high recall (0.937), while ResFinder maintained more balanced performance (precision 0.752–0.785, recall 0.793–0.807). Similarly, predictions for macrolides and lincosamides showed moderate concordance, erythromycin and clindamycin predictions showed only low concordance with phenotypic observations.

To test for the presence of known and potentially novel mechanisms of antimicrobial resistance, we constructed a pangenome (Supplementary Figure S8) and performed a pangenome-wide association study using Scoary [20] (Supplementary Table S4). The result of the bacterial GWAS identified known mechanisms of antimicrobial resistance. Genes such as *mecA* and *mecR1* were strongly associated with oxacillin and cefoxitin resistance. Similarly, the aminoglycoside resistance was determinated by *aacA-aphD* and *knt* for tobramycin; tetracycline resistance was shown to be mediated by efflux pump *tet(K)*; and the genes *ermC* and *msrA* were associated with resistance to macrolides and lincosamides (erythromycin, clindamycin).

However, non-canonical association were present for other drugs classes. For instance, fluoroquinolone resistance was associated with genes such as *hsdM* and *entC2*. Benzylpenicillin resistance associated pre-dominantly with *fcl_1*. Fusidic acid resistance showed associations primarily with transport-related genes (*natA, macB*, and *dppB*).

### Virulence and pathogenicity

The prevalence and distribution of toxin and virulence factors indicates regional variation (see Figure 1 and Supplementary Table S5). Panton-Valentine Leukocidin (PVL), a cytotoxin often associated with community-associated MRSA (CA-MRSA), was detected in 17% of isolates overall. However, its prevalence showed marked regional differences, being highest in Madinah (32.4%) and lowest in Hail (6.3%). Notably, PVL positivity was linked to specific lineages found predominantly in pilgrimage cities, including the internationally recognized CC1/ST772 (Bengal Bay clone).

Toxic shock syndrome toxin (TSST) was found in 10% of isolates and TSST rates showed an inverse geographic distribution pattern, ranging from 0% in Jazan to 26% in Hail, with intermediate rates in pilgrimage cities: Jeddah (8%), Makkah (8%), and Madinah (13%). Western cities (Jeddah, Madina, and Makkah) harbored the majority of Novel (ACME)-positive isolates.

Core virulence factors displayed widespread but distinctive distribution patterns across our isolates. Adhesion factors showed varying prevalence: clumping factors (clfA, clfB) appeared in over 95% of isolates, while the collagen binding protein (cna) occurred exclusively in CC5. The majority of isolates (>90%) carried immune evasion genes (*sak, sbi, scn*) and biofilm formation genes (icaABCD cluster). Among toxin genes, *sea* dominated with 30% prevalence, followed by other enterotoxins (*seb, sec, sed, seh*).

## Discussion

### Impact of mass gatherings on MRSA diversity and resistance patterns

This study described the genetic epidemiology of *S. aureus* across Saudi Arabia, a country with over 32 million inhabitants of which 41% are immigrants [11]. As of 2023, Saudi Arabia hosts over 2 million pilgrims for Hajj and 27 million pilgrims for Umrah annually [12]. While previous work has investigated the effect of mass gatherings and pilgrimage on antimicrobial resistance in Saudi Arabia [28], our integrated genotype–phenotype dataset reveals several distinct patterns associated with mass gathering activities.

The impact of mass gatherings on MRSA epidemiology is particularly evident in cities along the Hajj and Umrah routes. Jeddah, Makkah, and Madina showed significantly higher genetic diversity and unique strain patterns compared to other regions, with Jeddah, the main entry point for pilgrims, exhibiting the highest number of novel sequence types. This pattern aligns with observations from other Gulf regions, where mass gatherings coincide with diverse strain distributions [48, 19]. This diversity also includes a prevalence of internationally recognized strains, such as the PVL-positive Bengal Bay clone (CC1-ST772) and the South Pacific clone (CC30-ST30), which are found in pilgrimage cities, indicating a link between pilgrimage routes and the importation of globally circulating lineages [16].

This findings presented suggest that mass gatherings influence not only genetic diversity but also antimicrobial resistance patterns. Pilgrimage cities showed significantly higher overall resistance rates compared to other locations, suggesting these sites may serve as hotspots for resistance transmission. This is particularly evident in the distribution of SCC*mec* types, which suggests a shift from hospital-associated to community-associated MRSA, reflecting global trends [24, 37]. The concentrated presence of ACME-positive isolates in western pilgrimage cities further supports the role of international travel in strain dissemination.

The successful persistence of these strains in mass gathering-associated regions appears to be facilitated by multiple factors. The high prevalence of adaptive features, including biofilm formation genes and immune evasion clusters, alongside various resistance mechanisms, may enable these strains to be established in these high-traffic areas. These findings extend previous work on the impact of mass gatherings on bacterial populations [1], demonstrating how such events can facilitate the concurrent spread of both antimicrobial resistance determinants and virulence factors.

This unique demographic pattern and its influence on MRSA evolution and clonal expansion [34] presents both challenges and opportunities for public health interventions. Our findings suggest that large travel hub cities may serve as entry points for emerging strains and resistance patterns, potentially allowing early detection of concerning variants before they achieve wider distribution.

CC97 is detected as MSSA, and MRSA carrying SCCmecV associated with FusC gene. Newly identified ST8637 in Madinah is an indicator of active expansion of this linage and successful distribution in the community. The presence of CC97 in clinical samples indicates successful establishment in hospitals. More investigations are needed to understand the genetic basis of this adaptation and its implications for public health, especially as it ranks second in presence across the regions.

Interestingly, a significant increase in the proportion of novel sequence type assignments in Jeddah (9/14) was observed, and this the main entry point for international travelers to Hajj and Umrah, and provides a further indication that pilgrimage is one of the main drivers of drug resistance and diversity in Saudi Arabia.

### Limitations of Genomic Prediction for Antimicrobial Resistance

These findings highlight the complex relationship between genotype and phenotype in antimicrobial resistance prediction. While genomic prediction tools showed high precision for certain antibiotics (over 0.98 for beta-lactams and fusidic acid), they performed poorly for other drug classes, particularly aminoglyco-sides. This variable performance reflects both the complexity of resistance mechanisms and the current limitations of prediction methods.

The GWAS analysis revealed both expected and unexpected genetic associations. While we confirmed established resistance determinants such as *mecA_1* and *mecR1* for beta-lactams, and *ermC* and *msr(A)* for macrolide resistance — we also identified several unexpected associations requiring careful interpretation. For instance, fluoroquinolone resistance showed associations with *hsdM* and *entC2*, likely representing indirect relationships rather than causative mechanisms. Similarly, the association of fusidic acid resistance with transport-related genes (*natA, macB, dppB*) rather than expected target site mutations suggests potential limitations in our GWAS approach for detecting point mutations.

This discrepancy between genetic markers and phenotypic resistance highlights several challenges. In particular, resistance mechanisms may involve complex genetic interactions that are not captured by current prediction tools. These may include environmental factors and gene expression regulation which can contribute to resistance phenotypes independently of genetic markers. The convergence of diverse international strains in our study setting may facilitate the exchange of resistance determinants through mobile genetic elements, complicating genotype–phenotype relationships.

Given these limitations, we would suggest maintaining phenotypic testing alongside genomic surveillance for effective clinical decision-making [15]. Such a dual approach is especially crucial in regions experiencing high levels of international travel and strain diversity, where rapid evolution and transmission of resistance mechanisms may occur. Future improvements in prediction accuracy will likely require integration of multiple data types, including transcriptomics and proteomics, to better capture the complex determinants of antimicrobial resistance.

### Data Availability and Reproducibility

A key contribution of this study is the comprehensive implementation of FAIR (Findable, Accessible, Interoperable, and Reusable) principles [54] in data sharing. We structured our metadata using the Resource Description Framework (RDF) [33], creating a semantically rich representation of both sequence data and phenotypic measurements. Our data model (Supplementary Figure S3) incorporates standard terminologies and ontologies including the NCBI Taxonomy [45] for species annotation, the ChEBI ontology [29] for antimicrobial agents, the NCI Thesaurus [26] and the GENEPIO ontology [27] for health status descriptions, and the Unit Ontology (UO) [25] for standardized units, enhancing interoperability with existing datasets.

This structured approach to data sharing serves multiple purposes. First, it enables direct integration with other surveillance datasets through standardized terminology and relationships. The use of controlled vocabularies for key metadata elements — such as specimen sources, host characteristics, and antimicrobial susceptibility measurements — facilitates automated data integration and comparative analyses across different studies and geographical regions.

Second, this dataset reported is particularly valuable for developing and validating machine learning approaches for antimicrobial resistance prediction. By providing paired genomic and phenotypic data in a standardized format, along with detailed host and environmental metadata, we enable researchers to develop prediction models that can account for contextual factors beyond purely genetic determinants, and may be more adapted to the complex demographics found among pilgrims. The comprehensive antimicrobial susceptibility data, including both MIC values and categorical interpretations, further provides rich training and validation data for such models.

The complete computational reproducibility of our analysis, implemented through a workflow in the Common Workflow Language (CWL) [8], ensures that other researchers can not only access our data but also reproduce and build upon our analytical methods. By making our workflows, source code, and execution environments publicly available, we facilitate easier adaptation of our methods to new datasets and research questions.

Implementation of FAIR principles extends beyond technical accessibility to practical reusability. The standardized metadata format allows researchers to integrate our findings with their local surveillance data, potentially revealing new patterns of MRSA transmission and evolution. Furthermore, our dataset can serve as a baseline for tracking changes in MRSA populations, particularly in the context of mass gatherings and international travel, where standardized data collection and sharing are crucial for effective surveillance.

### Future Work

Our findings as reported point to several key directions for future surveillance strategies, particularly focusing on early detection at pilgrimage entry points. The implementation of systematic sampling could be strategically enhanced by targeting major travel hubs. Specifically, wastewater surveillance at airports serving pilgrimage routes, particularly the international airports in Jeddah and Madinah, could provide early warning signals of emerging strains. Aircraft wastewater sampling from pilgrimage flights, which has proven effective for COVID-19 surveillance [50], could be adapted for MRSA monitoring, offering insights into strain importation patterns before pilgrims arrive at their destinations.

Such targeted approach could be complemented by environmental sampling at key congregation points along pilgrimage routes. Strategic sampling points could include ablution facilities in major mosques and shared sanitation facilities in pilgrim accommodations, providing a comprehensive picture of strain circulation during mass gatherings. Integration of these environmental samples with clinical surveillance would create a more complete understanding of MRSA transmission dynamics during mass gathering.

The application of long-read sequencing technologies to these samples could resolve complex genetic elements that are challenging to characterize with short-read data alone. This would be particularly valuable for understanding the structure of SCCmec cassettes and other mobile genetic elements that may be exchanged during mass gatherings.

Finally, establishing a real-time surveillance system that combines this strategic sampling with rapid genomic and phenotypic analysis would enable faster response to emerging resistant strains. Such a system could serve as a model for monitoring other pathogens in similar mass gathering settings, contributing to global pathogen surveillance efforts and early warning systems for emerging antimicrobial resistance.

## Conclusion

In conclusion this study demonstrates how mass gatherings and regional factors jointly shape bacterial population structures and antimicrobial resistance patterns at a national scale. While pilgrimage cities exemplify the intersection of global human mobility with pathogen evolution, the distinct patterns we observed in non-pilgrimage regions highlight the importance of local healthcare practices and potential livestock–human transmission routes in shaping MRSA populations. This comprehensive view, spanning both high-traffic pilgrimage sites and regional healthcare settings, provides evidence for a framework to understand pathogen dynamics in complex environments where international, community, and agricultural factors converge. Through standardized surveillance approaches and data sharing, our work establishes a foundation for monitoring antimicrobial resistance in the context of both global human mobility and regional One Health challenges, with implications for public health strategies across the Middle East and beyond.

## 1 Data Availability Statement

Sequencing data is available on the Sequence Read Archive (SRA) as a BioProject under accession number `PRJEB59751` The phenotypes for each samples are available on Zenodo under DOI `10.5281/zenodo.1425085` [36].

## 2 ETHICS APPROVAL

This work approved by the Institutional Biosafety and Bioethics Committee (IBEC) of King Abdullah University of Science and Technology under approval number `18IBEC14`.

