## supplemental material for "Genomic diversity and antimicrobial resistance of *Staphylococcus aureus* in Saudi Arabia: a nationwide study using whole-genome sequencing"

### Supplementary Material for: Genomic surveillance of Methicillin Resistant Staphylococcus aureus in Saudi Arabia

[1]Computational Bioscience Research Center, King Abdullah University of Science and Technology, KAUST, [2]SDAIA–KAUST Center of Excellence in Data Science and Artificial Intelligence, King Abdullah University of Science and Technology, KAUST, [3]Environmental Science and Engineering Program, Division of Biological and Environmental Science and Engineering, KAUST, [4]Wellness and Preventive Medicine Institute, Health Sector, King Abdulaziz City for Science and Technology (KACST), Riyadh, Saudi Arabia, [5]College of Applied Medical Sciences, Taibah University, Madinah, Saudi Arabia, [6]Department of Medical Laboratory Technology, College of Applied Sciences, Taibah University, Madinah, Saudi Arabia, [7]Department of biological sciences, college of

science, University of Jeddah, Jeddah, Saudi Arabia., [8]Vaccines and Immunotherapy Unit, King Fahd Medical Research Center, King Abdulaziz University, Jeddah 21589, Saudi Arabia, [9]Department of Medical Laboratory Technology, Faculty of Applied Medical Sciences, King Abdulaziz University, Jeddah 21589, Saudi Arabia, [10]Epidemiology department, public health Administration, king Abdullah Medical Complex, Jeddah 23816, Saudi Arabia, [11]Alnoor Specialist Hospital, Ministry of Health, Makkah, Saudi Arabia, [12]Infectious Diseases Research Department, King Abdullah International Medical Research Center (KAIMRC), Riyadh, Saudi Arabia., [13]King Abdullah International Medical Research Center (KAIMRC), [14]King Saud bin Abdulaziz University-Health Sciences Ministry of National Guard-Health Affairs (MNGHA), [15]Department of Pathology and Laboratory Medicine, King Abdulaziz Medical City (KAMC),, [16]Ministry of National Guard Health Affairs (MNGHA), Riyadh, Saudi Arabia., [17]World Health Organization, Geneva, Switzerland, [18]Medical Laboratory, King Abdulaziz Medical City (KAMC), Ministry of National Guard Health Affairs, Jeddah, Saudi Arabia, [19]Emerging and Epidemic Infectious Diseases Research Unit, Medical Research Center, Jazan University, Jazan 45142, Saudi Arabia, [20]Department of Medical Laboratories Technology, College of Applied Medical Sciences, Jazan University, Jazan, Saudi Arabia, [21]Department of Medicine,Faculty of Medicine, Jazan University, Jazan, Saudi Arabia, [22]Regional Laboratory & Central Blood Bank Jazan Health, [23]Saudi Public Health Authority, Vector-Borne Diseases Laboratory, Jazan 45142, Saudi Arabia, [24]BndrGene Medical Lab, Madinah, Saudi Arabia, [25]Faculty of Pharmaceutical Sciences, Department of Pharmacy, Tokyo University of Science, Noda, Chiba, Japan, [26]Department of pathology, college of Medicine, King Saud University and King Saud University Medical City, Riyadh, Saudi Arabia, [27]The National Center for Genomic Technology (NCGT), Life Science and Environment Research Institute, King Abdulaziz City for Science and Technology (KACST), Riyadh, Saudi Arabia, [28]Medical Microbiology Laboratory, Hera General Hospital, Makkah healthcare cluster, Makkah, Saudi Arabia., [29]KAUST Center of Excellence for Smart Health (KCSH), King Abdullah University of Science and Technology, 4700 KAUST, Thuwal 23955, Saudi Arabia, [30]KAUST Center of Excellence for Generative AI, King Abdullah University of Science and Technology, 4700 KAUST, Thuwal 23955, Saudi Arabia, [31]Biothreat Response Department, Public Health Laboratory, the Saudi Public Health Authority, [32]Computer, Electrical and Mathematical Sciences & Engineering (CEMSE) Division, King Abdullah University of Science and Technology, King Abdullah University of Science and Technology, 4700 KAUST, Thuwal, Saudi Arabia., [33]Biological and Environmental Sciences & Engineering (BESE) Division, King Abdullah University of Science and Technology, King Abdullah University of Science and Technology, 4700 KAUST, Thuwal, Saudi Arabia, [34]Advanced Diagnostics and Therapeutics Institute, Health Sector, King Abdulaziz City for Science and Technology (KACST), Riyadh, Saudi

Arabia, [35]UCD-Centre for Food Safety, University College Dublin, Belfield, Dublin D04 N2E5, Ireland, [36]nstitute for Global Food Security (IGFS), The Queen's University of Belfast, 19 Chlorine Gardens, Belfast BT9 5DL, Northern Ireland, United Kingdom, [37]King Faisal Specialist Hospital & Research Centre- Madina

May 10, 2025

### Supplementary Figures

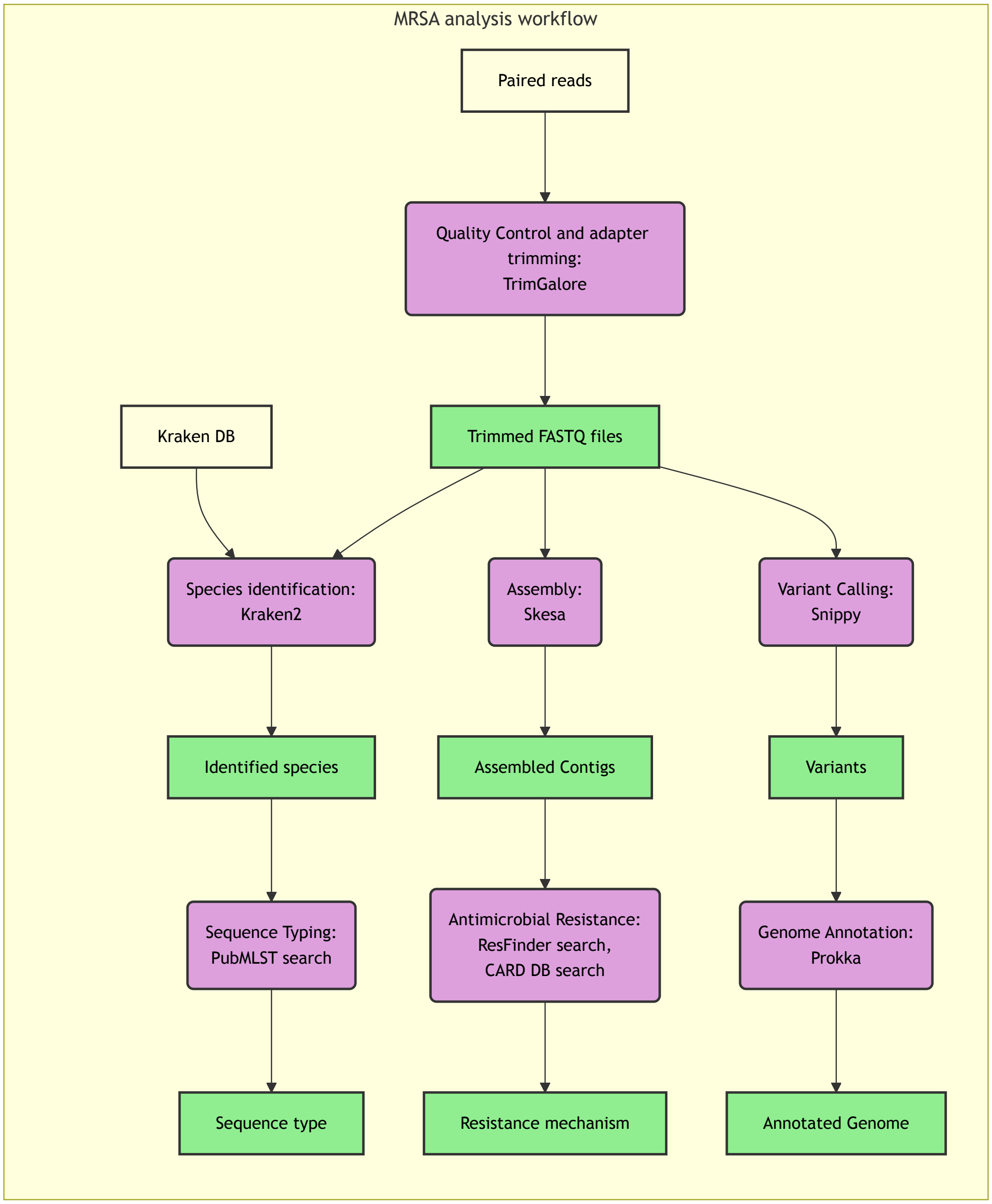


Figure S1: This diagram outlines the key steps performed on the raw sequencing reads for each sample, including quality control and adapter trimming (TrimGalore), taxonomic identification (Kraken2), genome assembly (SKESA), variant calling (Snippy), MLST typing (ABRicate/PubMLST), antimicrobial resistance gene detection (ABRicate/ResFinder/CARD), and genome annotation (Prokka). The workflow is designed to be executed in a CWL-compliant environment, allowing for reproducible and scalable analysis of genomic data.

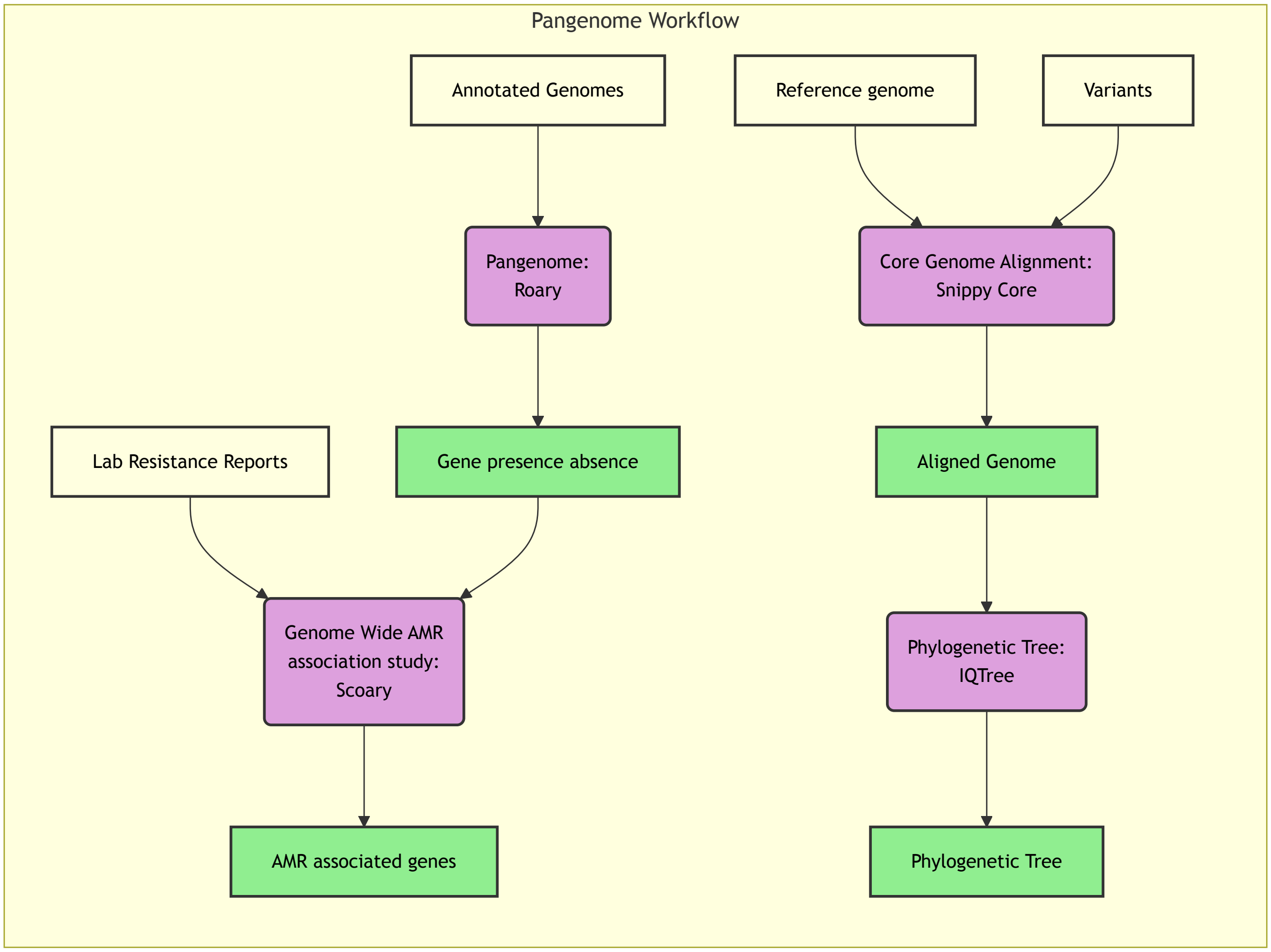


Figure S2: This diagram illustrates the steps following initial processing (Figure S1). Indicates pangenome construction from annotated genomes (Roary), core genome alignment generation (Snippy Core), phylogenetic tree inference (IQTree), and the pangenome-wide association study (Scoary) correlating gene presence/absence with phenotypic resistance data. [5]This workflow was also implemented using CWL-compliant environment.

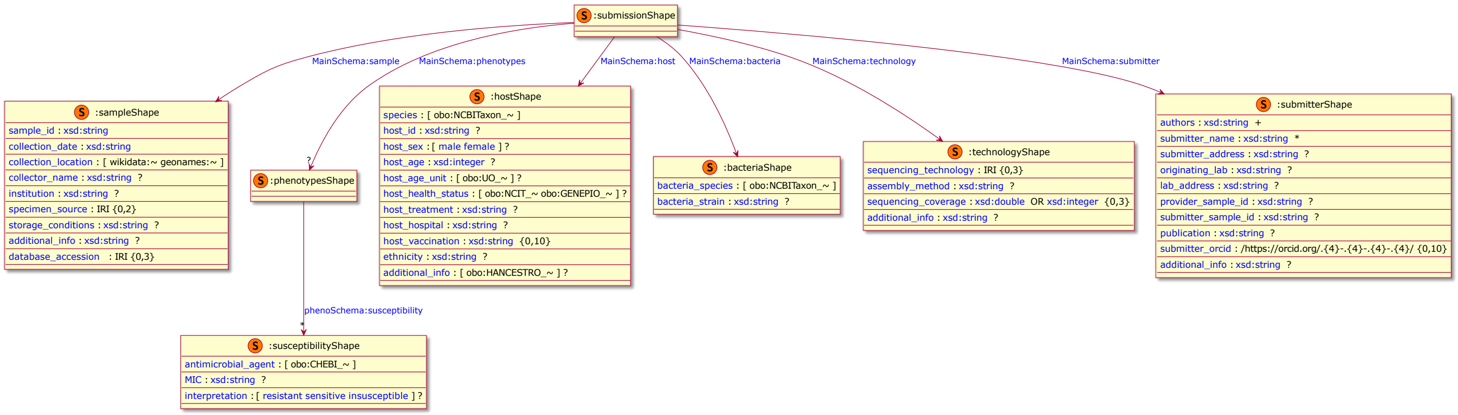


Figure S3: Metadata RDF data model. ? indicates an optional field, + indicates one or more values, ⋆ indicates zero or more values. This schema illustrates the structure used to represent sample, host, bacterial, sequencing, submitter, and phenotypic susceptibility information in RDF format. It leverages standard ontologies to ensure interoperability and adherence to FAIR data principles, facilitating data integration and reuse.

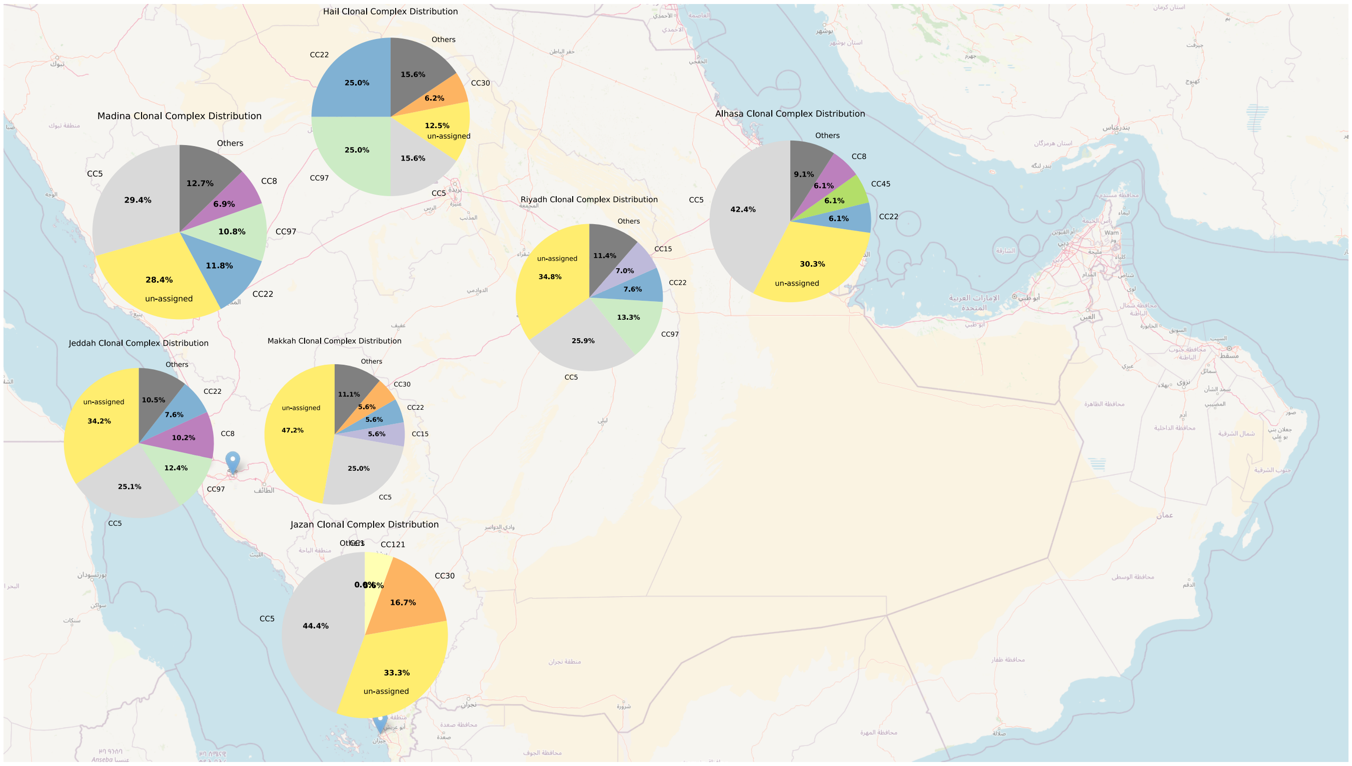


Figure S4: Geographic distribution of major *S. aureus* Clonal Complexes (CCs) across Saudi Arabia. The map represents the prevalence of the top five most abundant CCs (CC5, CC22, CC97, CC30, CC8) across regions. Pie charts indicate the relative proportion of these major CCs, visually demonstrating regional differences in lineage composition.

### Supplementary Tables

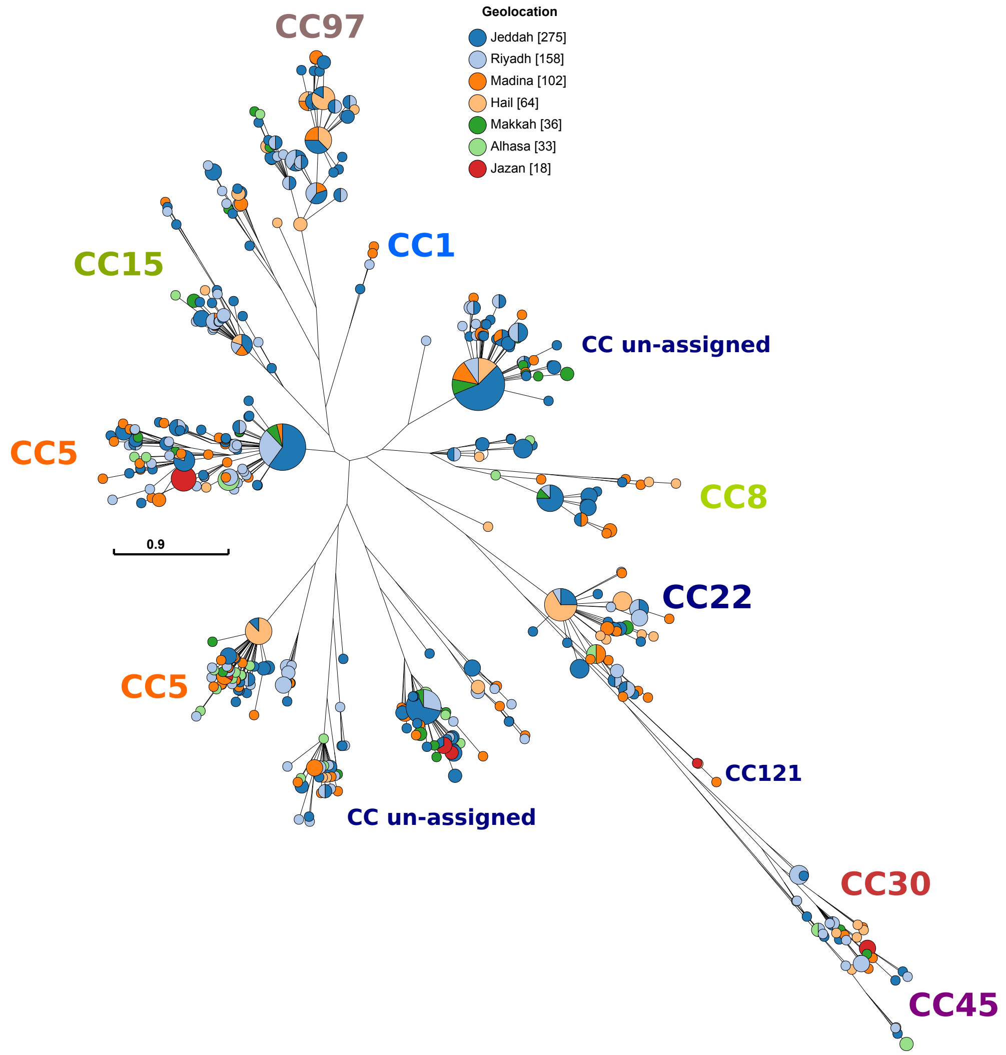


Figure S5: Minimum Spanning Tree indicating genetic relatedness and geographic structure. Network visualization based on core genome SNP distances, showing the genetic relationships between isolates. Each node represents isolates, sized frequency, and colored by geographic region of origin. Major Clonal Complexes (CCs) are labelled, demonstrating the genetic clustering of related strains and how geographic origins are distributed within and across these major lineages or (CC).

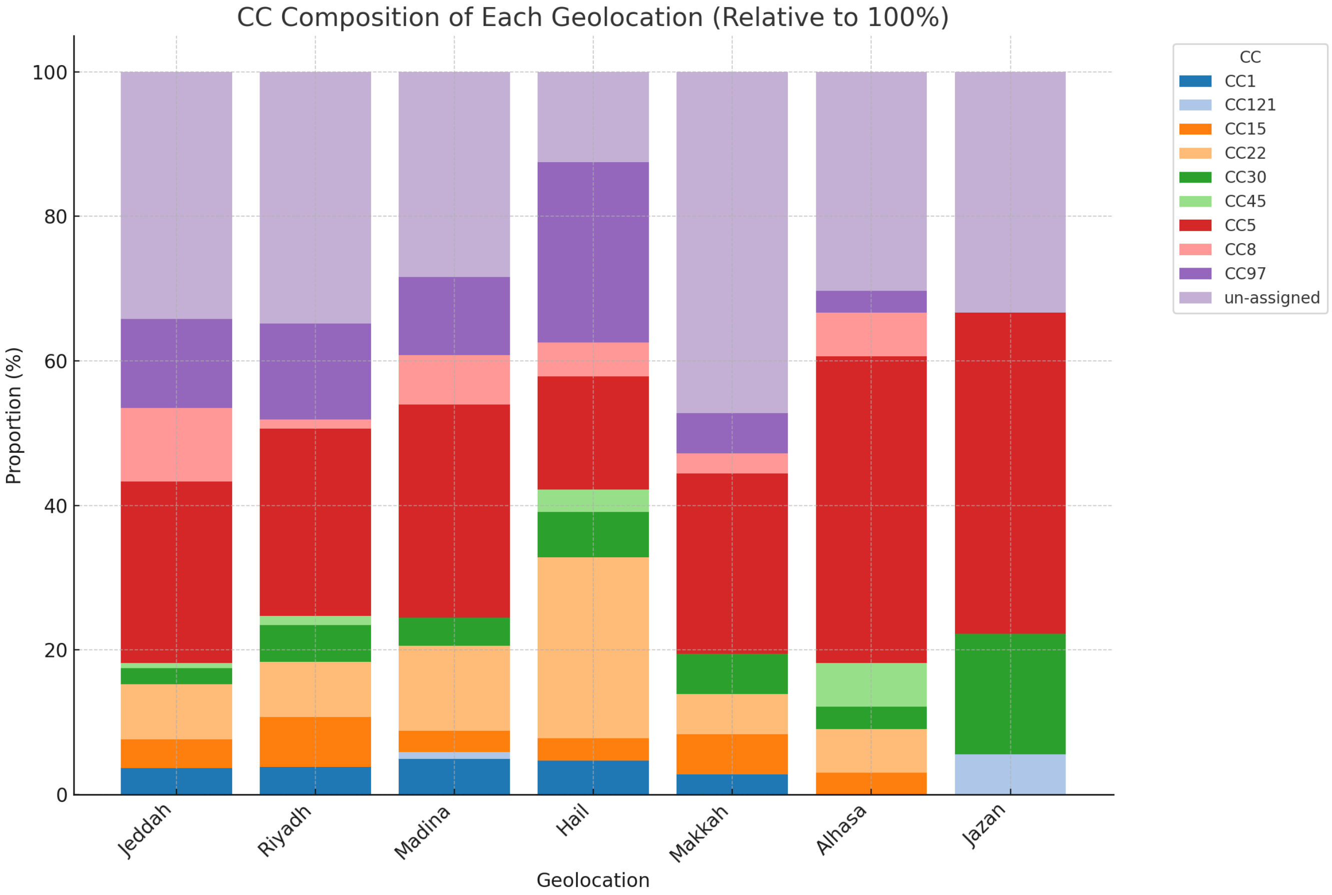


Figure S6: Relative abundance of Clonal Complexes (CC) for each region based on the samples size per region. Stacked bar chart indicates high prevalence of CC5, CC97, CC22, CC30, CC15, and unassigned (CC).

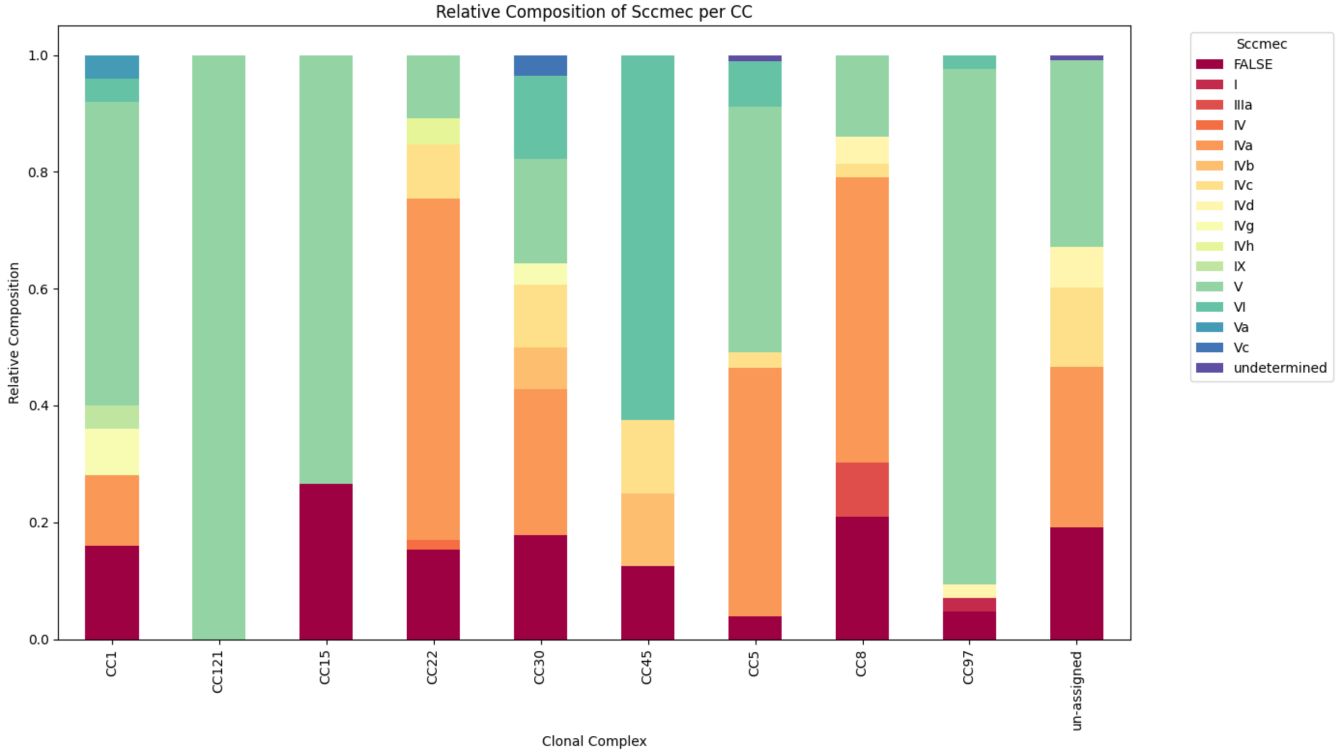


Figure S7: Stacked bar chart showing the distribution of different SCCmec types. The composition of Sccmec elements relative to clonal complex. The figure demonstrates lineage-specific associations, such as the high prevalence of SCCmec types IVa and V indicating CA-MRSA prevalence (CC5, CC22).

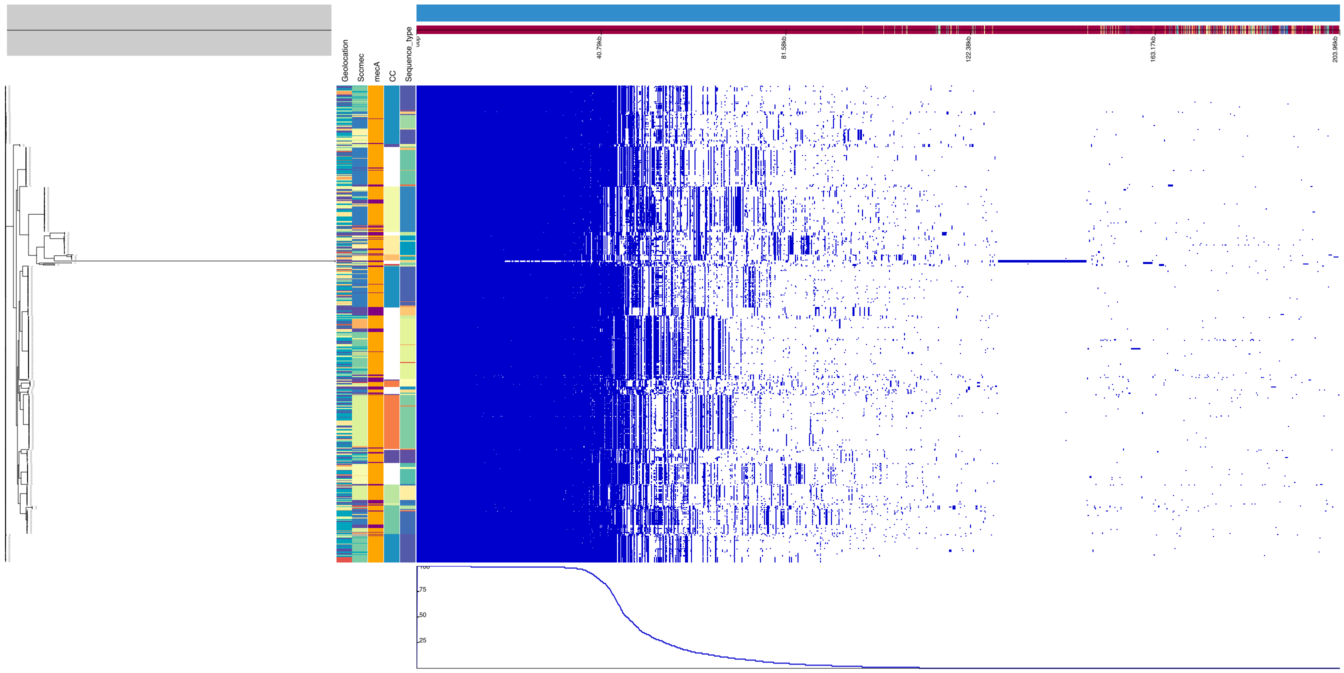


Figure S8: Visualization of the *S. aureus* pangenome matrix. Heatmap representing the presence (blue) and absence (white) of genes across the 686 isolates. Rows correspond to isolates (clustered by phylogeny on the left), and columns represent genes present in the pangenome (ordered by clustering).

| **Region** | **No. of Samples (Included in this Study)** |
|---|---|
| AlHasaa (East) | 37 (33) |
| Hail (North) | 69 (64) |
| Jazan (South) | 33 (18) |
| Jeddah (West) | 373 (275) |
| Madina (West) | 121 (102) |
| Makkah (West) | 38 (36) |
| Riyadh (Central) | 178 (158) |

Table S1: Samples counts per city/region, with the total count and included samples post quality check in parenthesis.

| Antibiotic | Concentration ($\mu$g/ml) |
|---|---|
| Benzylpenicillin | 0.125, 0.25, 1 |
| Oxacillin | 0.5, 1, 2 |
| Gentamicin | 8, 16, 64 |
| Tobramycin | 16, 32, 64 |
| Levofloxacin | 0.25, 2, 8 |
| Moxifloxacin | 0.25, 2, 8 |
| Erythromycin | 0.25, 0.5, 2 |
| Clindamycin | 0.5, 1, 2 |
| Linezolid | 0.5, 1, 2 |
| Teicoplanin | 1, 4, 8, 16 |
| Vancomycin | 1, 4, 8, 16 |
| Tetracycline | 0.5, 1, 2 |
| Tigecycline | 0.25, 0.5, 1 |
| Fosfomycin | 8, 32 |
| Nitrofurantoin | 16, 32, 64 |
| Fusidic acid | 0.5, 1, 4 |
| Mupirocin | 1 |
| Rifampicin | 0.25, 0.5, 2 |
| Trimethoprim/Sulfamethoxazole | 8/152, 16/304, 32/608 |

Table S2: Antimicrobial agents and their concentrations tested by the AST-P580 card

| ST | Location | SCCmec, SPA Type | Comments |
|---|---|---|---|
| ST8633 | Riyadh | SCCmecIVa-t304 | |
| ST8635 | Madina | Unresolved SCCmec profile | |
| ST8636 | Madina | SCCmecV-t3841 | CC97 |
| ST8638 | Alhassa | SCCmecIVa-t304 | CC5 |
| ST8639 | Jeddah | MSSA SPA t3341 | SLV of ST-88 |
| ST8641 | | SCCmecV | CC15, SLV of ST-1535 |
| ST8642 | Jeddah | MSSA-t008 | PVL+, ACME+, CC8 |
| ST8643 | Jeddah | SCCmecV-t3841 | SLV of ST-672 |
| ST8644 | Jeddah | SCCmecV | |
| ST8645 | Jeddah | SCCmecV-t311 | CC5 |
| ST8646 | Jeddah | MSSA | CC8 |
| ST8647 | Jeddah | SCCmecV-t991 | |
| ST8648 | Jeddah | SCCmecIVc | CC8 |
| ST8649 | Jeddah | SCCmecIVa-t2453 | |

Table S3: Summary of newly assigned ST types, locations, and SCCmec/SPAtype profiles in Saudi Arabia.

| Drug | List of Significant Genes |
|---|---|
| Moxifloxacin | *hsdM*, *entC2*, *lpl2_4*, *ssl7_1*, *entS_2*, *nikC*, *fnbB*, *hsdM_2*, *gtaB*, *fhuD_1* |
| Gentamicin | *aacA-aphD*, *rarD*, *adhR*, *lpl2_6*, *farB_1*, *isp*, *salL*, *femA_1*, *adhE*, *lpl2_5* |
| Benzylpenicillin | *fcl_1* |
| Cefoxitin | *mecA_1*, *ugpQ*, *mvaS_1*, *mecR1*, *ssl1* |
| Levofloxacin | *hsdM*, *entS_2*, *entC2*, *hsdM_2*, *cadC*, *fosB*, *fhuD_1*, *fhuD*, *gntR*, *ureC* |
| Tobramycin | *aacA-aphD*, *knt*, *lpl2_5*, *rarD*, *salL*, *xerC_1*, *sigS*, *bglA*, *adhR*, *gntR* |
| Oxacillin | *mecA_1*, *ugpQ*, *mvaS_1*, *mecR1*, *prmC_1* |
| Trimethoprim-Sulfamethoxazole | *entD*, *blaZ*, *xerC_4*, *pepT_2*, *thyA_1*, *lpl2_2*, *essG_1*, *manR_1*, *yjdF*, *entB* |
| Tetracycline | *tet(K)*, *pre*, *linA*, *cadC*, *lagD*, *knt*, *aphA*, *entE*, *sirC*, *satA* |
| Erythromycin | *ermC*, *msr(A)*, *bcrA_3*, *graS_3*, *ble*, *bcrB* |
| Fusidic Acid | *yoaA*, *wbnH*, *yknY*, *arsB*, *natA*, *dppB*, *arsC*, *gsiA*, *gsiC_1*, *macB* |
| Clindamycin | *ermC* |

Table S4: Top significant genes (Bonferonni corrected p-value $< 0.05$) associated with resistance to various antibiotics identified through GWAS analysis using Scoary v1.6.16. The list includes genes significantly correlated with resistance to each antibiotic tested.

| Gene | FALSE | TRUE | %FALSE | %TRUE |
|---|---|---|---|---|
| sea | 474 | 212 | 69.09 | 30.90 |
| seb | 593 | 93 | 86.44 | 13.55 |
| sec | 660 | 26 | 96.20 | 3.79 |
| sed | 666 | 20 | 97.08 | 2.91 |
| seh | 638 | 48 | 93.00 | 6.99 |
| selk | 635 | 51 | 92.56 | 7.43 |
| sell | 660 | 26 | 96.20 | 3.79 |
| selq | 636 | 50 | 92.71 | 7.28 |
| TSST | 614 | 72 | 89.50 | 10.4 |
| ACME | 660 | 26 | 96.20 | 3.79 |

Table S5: Toxin and virulence genes identified in our dataset.
